# Gut Virome Diversity and Health Status in Rescued Pangolins: Metaviromics Reveals Parvovirus as a Key Virus Associated with Disease

**DOI:** 10.64898/2026.07.16.738944

**Authors:** Wenjing Jiao, Zhiliao Zeng, Xinyuan Hu, Jianfeng Deng, Jinping Chen

## Abstract

**Background:** Pangolins are critically endangered mammals that suffer from high rates of gastrointestinal disease during captivity, yet the role of the gut virome in their health remains unexplored. This study presents the first comprehensive characterization of the gut DNA virome in Malayan (*Manis javanica*) and Chinese (*M. pentadactyla*) pangolins across different health states.

**Results:** Metaviromic sequencing of 16 fecal samples from healthy, diarrheal, pneumonic, free-ranging, and deceased pangolins generated 7.2–11.8 Gb clean data per sample. A total of 12 viral phyla, 26 families, 219 genera, and 1,132 species were identified. Caudovirales phages (Siphoviridae, Myoviridae, and Podoviridae) dominated the gut virome of healthy individuals, with phage content exceeding 90% in most healthy samples. However, diseased and deceased individuals exhibited a significant reduction in phage proportion (94.5% vs. 53.3%, P = 0.02), accompanied by a dramatic increase in eukaryotic viruses—particularly Parvoviridae, which accounted for 66% of the virome in deceased Malayan pangolins. Iridoviridae (38%) and Polydnaviridae (34%) dominated in deceased Chinese pangolins. SIMPER analysis identified Parvoviridae as the primary contributor to virome dissimilarity between healthy and diseased Malayan pangolins, whereas Iridoviridae and Polydnaviridae were the key contributors for Chinese pangolins. LEfSe analysis revealed 12 biomarker viruses in free-ranging pangolins (predominantly Staphylococcus and Escherichia phages), six in healthy Chinese pangolins (predominantly Streptococcus phages and Lactobacillus viruses), and six in deceased Malayan pangolins (predominantly vertebrate viruses). KEGG functional annotation revealed that genes related to DNA replication, repair, and recombination were the most abundant, suggesting frequent genomic recombination within the pangolin gut virome.

**Conclusions:** The gut virome of pangolins is closely associated with health status. Disease induces a shift from a phage-dominated to a eukaryotic virus–dominated virome, with Parvoviridae and Iridoviridae emerging as candidate pathogenic viruses in Malayan and Chinese pangolins, respectively. These findings provide molecular evidence for virome monitoring in pangolin conservation and highlight the need for targeted surveillance of

**Importance:** Pangolins are the most trafficked mammals in the world, and all eight species are critically endangered. During rescue and captive care, these animals suffer from high rates of gastrointestinal disease, yet the role of viruses in their gut health has never been studied. This research provides the first comprehensive map of the gut virus community in pangolins, revealing that healthy individuals carry a virus community dominated by beneficial bacterial viruses called phages, while sick and dying animals show a dramatic shift toward disease-causing viruses. Notably, parvoviruses emerge as potential pathogens in Malayan pangolins, while iridoviruses dominate in deceased Chinese pangolins. These findings offer a new tool for monitoring pangolin health in rescue centers: by tracking the balance between beneficial phages and harmful viruses, veterinarians may be able to detect illness earlier and improve survival rates. The study also highlights the importance of environmental exposure for maintaining a healthy gut virus community, supporting the creation of semi-natural enclosures for rescued pangolins. Beyond conservation, this work contributes to our understanding of how viruses move between wildlife and humans, which is critical for preventing future disease outbreaks.

## Introduction

Mammalian viromes encompass viruses infecting eukaryotic cells (eukaryotic virome), bacteriophages infecting bacteria (phage virome), and viruses infecting archaea (archaeal virome) (1). Viral-derived genetic elements integrated into host chromosomes can alter host gene expression and protein production, or even generate infectious virions (2). Unlike bacterial infections, which are typically localized, viral infections can persist systemically across nervous, hematopoietic, and vascular tissues, exerting sustained effects on host physiology (3, 4, 5). Mammalian viromes evolve continuously through rapid mutation, environmental exposure, and cross-species transmission (6, 7), making their study critical for understanding emerging infectious diseases.

Recent advances in viral metagenomics (metaviromics) have enabled culture-independent characterization of entire viral communities from environmental and clinical samples (8). This approach has revealed that the gut virome plays important roles beyond pathogenesis—shaping immune development, modulating bacterial community structure through phage predation, and influencing host metabolism (9, 10). In humans, virome dysbiosis has been associated with inflammatory bowel disease (9), COVID-19 (11, 12), and Clostridioides difficile infection (13). However, the gut viromes of wildlife species, particularly endangered mammals, remain largely uncharacterized.

Pangolins (Order Pholidota) are among the most trafficked mammals globally, with all eight species listed on CITES Appendix I (14). In China, both the Malayan pangolin (Manis javanica) and the Chinese pangolin (M. pentadactyla) are classified as National Class I Protected Species. During rescue and captive management, pangolins are highly susceptible to gastrointestinal infections, diarrhea, and parasitic infestations, which are the primary causes of mortality (15, 16, 17, 18). While previous studies have examined the gut bacteriome of pangolins (19, 20, 21, 22), the gut virome—particularly its relationship with health status—has not been investigated.

Pangolins have also attracted considerable attention as potential intermediate hosts for SARS-CoV-2 (18, 23, 24), highlighting the public health significance of characterizing their viral communities. Understanding the baseline virome composition of pangolins under different health conditions is essential for both conservation and zoonotic disease surveillance.

In this study, we performed metaviromic sequencing on fecal samples from 16 pangolins representing six health states (healthy Malayan, healthy Chinese, diarrheal Malayan, pneumonic Malayan, free-ranging Malayan, and deceased individuals). Our objectives were to: (1) characterize the overall diversity and composition of the pangolin gut DNA virome; (2) identify health-associated shifts in virome structure; (3) determine key viral taxa contributing to virome dysbiosis in diseased individuals; and (4) assess the functional potential of the pangolin gut virome.

## Results

Sequencing quality and data overview. After quality filtering, 16 samples yielded 7.2–11.8 Gb clean data, with clean read percentages ranging from 69.69% to 82.80% (Table 2). Following host removal, 24,468,296–34,468,052 non-host reads were obtained per sample. Host removal efficiency was high for most samples (81.16–99.98%), with the exception of two deceased individuals: GZM4 (46.11%) and GZM7 (84.51%), which contained substantially higher proportions of host DNA.

**Table 1.**
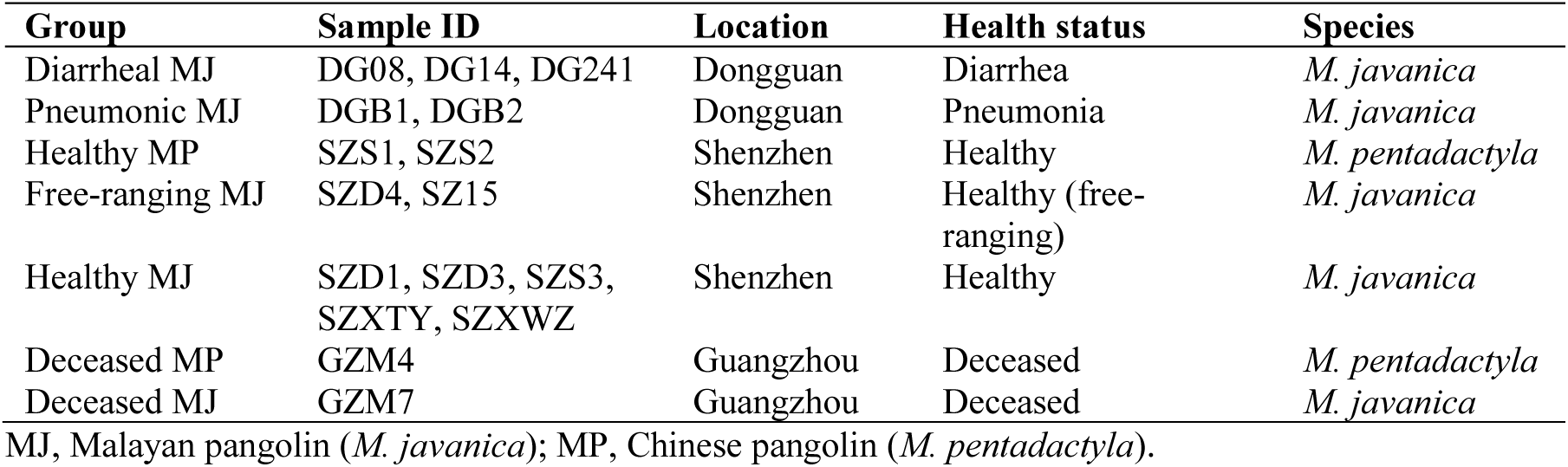
Sample information.

| Group | Sample ID | Location | Health status | Species |
| --- | --- | --- | --- | --- |
| Diarrheal MJ | DG08, DG14, DG241 | Dongguan | Diarrhea | *M. javanica* |
| Pneumonic MJ | DGB1, DGB2 | Dongguan | Pneumonia | *M. javanica* |
| Healthy MP | SZS1, SZS2 | Shenzhen | Healthy | *M. pentadactyla* |
| Free-ranging MJ | SZD4, SZ15 | Shenzhen | Healthy (free-ranging) | *M. javanica* |
| Healthy MJ | SZD1, SZD3, SZS3, SZXTY, SZXWZ | Shenzhen | Healthy | *M. javanica* |
| Deceased MP | GZM4 | Guangzhou | Deceased | *M. pentadactyla* |
| Deceased MJ | GZM7 | Guangzhou | Deceased | *M. javanica* |

MJ, Malayan pangolin (*M. javanica*); MP, Chinese pangolin (*M. pentadactyla*).

**Table 2.** Sequencing quality and host removal statistics.

| Sample | Raw_base(G) | Clean_base(G) | Clean_reads(PE) | Clean % | Non-host % |
| --- | --- | --- | --- | --- | --- |
| DG08 | 12.5 | 10.2 | 34483687 | 82.8 | 99.95 |
| DG14 | 11.4 | 9.4 | 31610340 | 83 | 99.98 |
| DG241 | 9.9 | 7.2 | 24473251 | 74.29 | 99.98 |
| DGB1 | 15.4 | 9.4 | 35759701 | 69.69 | 93.63 |
| DGB2 | 10.9 | 8.9 | 30236839 | 82.99 | 99.98 |
| GZM4 | 15.3 | 11.8 | 41160932 | 80.72 | 46.11 |
| GZM7 | 15.9 | 10.6 | 40331703 | 76.06 | 84.51 |
| SZ15 | 11.2 | 8.7 | 29620559 | 79.61 | 99.88 |
| SZD1 | 10.8 | 8.9 | 29975349 | 83.5 | 99.78 |
| SZD1.B | 11.6 | 9.2 | 31190587 | 80.36 | 98.08 |
| SZD3 | 9.9 | 7.5 | 25374205 | 76.59 | 99.94 |
| SZD4 | 11.4 | 9.3 | 31625667 | 83.2 | 99.04 |
| SZS1 | 11.4 | 9.1 | 31256270 | 82.08 | 98.27 |
| SZS2 | 12.6 | 10 | 33755938 | 80.13 | 98.44 |
| SZS3 | 11.3 | 9 | 30451582 | 80.66 | 99.98 |
| SZXTY | 11.5 | 9.2 | 31497724 | 82.31 | 81.16 |
| SZXWZ | 11.1 | 8.4 | 28655938 | 77.46 | 88.95 |

Phage content differs significantly between healthy and diseased pangolins. Viral category analysis revealed that phages constituted 14.23–99.96% of the total viral content across all samples (Table 3). Most healthy pangolin samples had phage content exceeding 90%, whereas diseased and deceased individuals showed markedly reduced phage proportions. One-way ANOVA confirmed a significant difference in phage content between healthy (mean = 94.5%) and diseased/deceased (mean = 53.3%) groups (P = 0.02). Notably, two diarrheal samples (DG08: 18.02% phages; DG241: 24.09% phages) and one deceased sample (GZM7: 14.23% phages) had dramatically lower phage proportions, suggesting that eukaryotic viral expansion accompanies disease onset.

**Table 3.** Viral category statistics.

| Sample | Other_virus(%) | Phages (%) |
| --- | --- | --- |
| DG08 | 81.98 | 18.02 |
| DG14 | 0.2 | 99.8 |
| DG241 | 75.91 | 24.09 |
| DGB1 | 25.76 | 74.24 |
| DGB2 | 15.05 | 84.95 |
| GZM4 | 41.93 | 58.07 |
| GZM7 | 85.77 | 14.23 |
| SZ15 | 21.85 | 78.15 |
| SZD1 | 2.83 | 97.17 |
| SZD3 | 0.04 | 99.96 |
| SZD4 | 1.04 | 98.96 |
| SZS1 | 2.76 | 97.24 |
| SZS2 | 6.86 | 93.14 |
| SZS3 | 2.76 | 97.24 |
| SZXTY | 1.88 | 98.12 |
| SZXWZ | 3.79 | 96.21 |

De novo assembly and viral annotation. Assembly of non-host reads produced 4,037–22,756 contigs per sample, with maximum lengths of 23,238–272,992 bp and N50 values of 479–2,947 (GC content: 35.25–56.01%). The majority of contigs (171,184) were shorter than 1,000 bp, while 1,983 contigs exceeded 10,000 bp.

Viral annotation identified a total of 12 phyla, 26 families, 219 genera, and 1,132 species across all samples (Table 4). The most frequently annotated viral families belonged to the order Caudovirales: Siphoviridae (2,338 contigs), Myoviridae (880 contigs), and Podoviridae (462 contigs). Among non-phage families, Parvoviridae (256 contigs), Genomoviridae (255 contigs), Microviridae (220 contigs), Iridoviridae (109 contigs), and Polydnaviridae (104 contigs) were the most represented (Table 5).

**Table 4.**
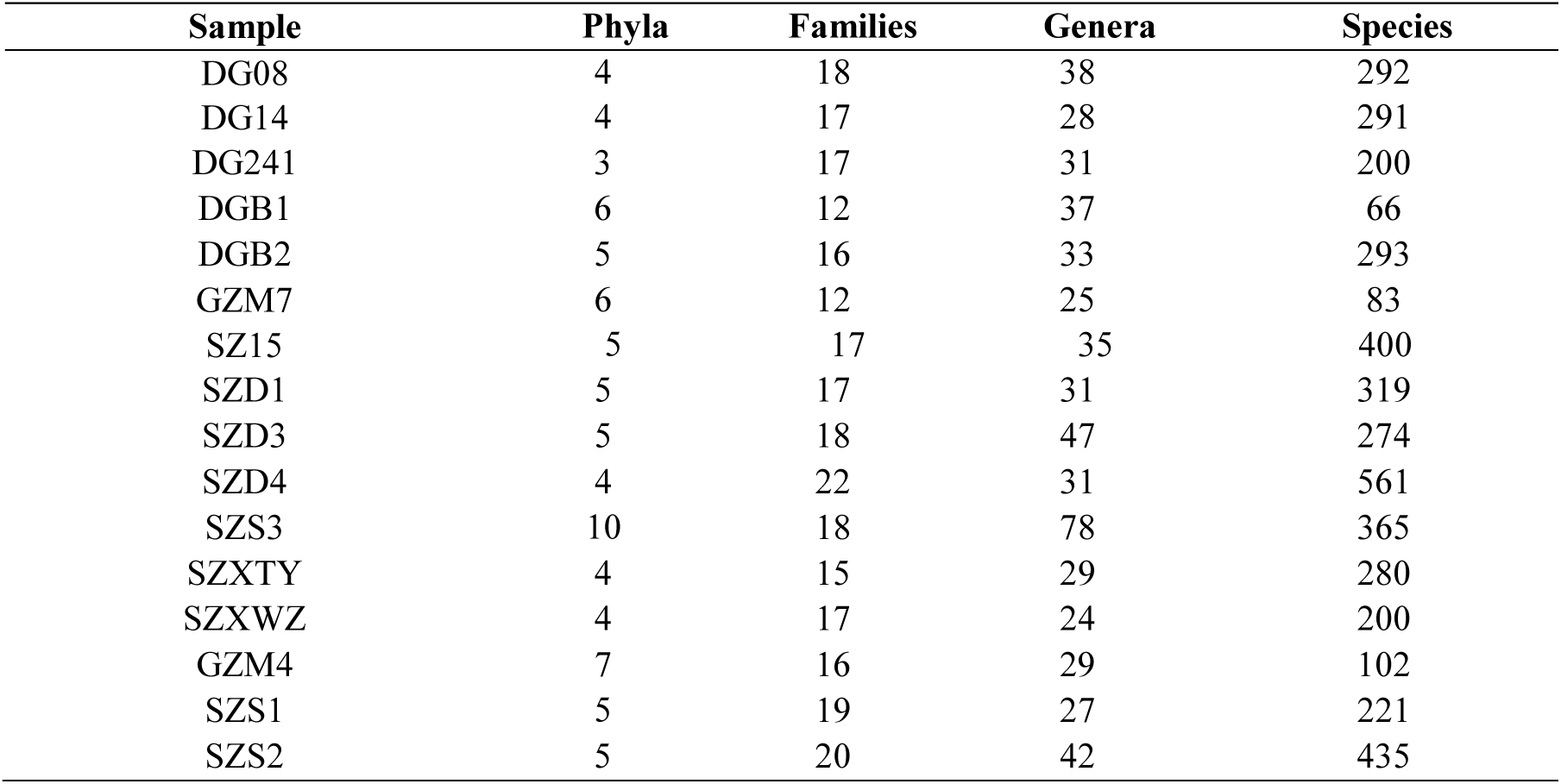
Viral taxonomic statistics per sample.

| Sample | Phyla | Families | Genera | Species |
| --- | --- | --- | --- | --- |
| DG08 | 4 | 18 | 38 | 292 |
| DG14 | 4 | 17 | 28 | 291 |
| DG241 | 3 | 17 | 31 | 200 |
| DGB1 | 6 | 12 | 37 | 66 |
| DGB2 | 5 | 16 | 33 | 293 |
| GZM7 | 6 | 12 | 25 | 83 |
| SZ15 | 5 | 17 | 35 | 400 |
| SZD1 | 5 | 17 | 31 | 319 |
| SZD3 | 5 | 18 | 47 | 274 |
| SZD4 | 4 | 22 | 31 | 561 |
| SZS3 | 10 | 18 | 78 | 365 |
| SZXTY | 4 | 15 | 29 | 280 |
| SZXWZ | 4 | 17 | 24 | 200 |
| GZM4 | 7 | 16 | 29 | 102 |
| SZS1 | 5 | 19 | 27 | 221 |
| SZS2 | 5 | 20 | 42 | 435 |

**Table 5.**
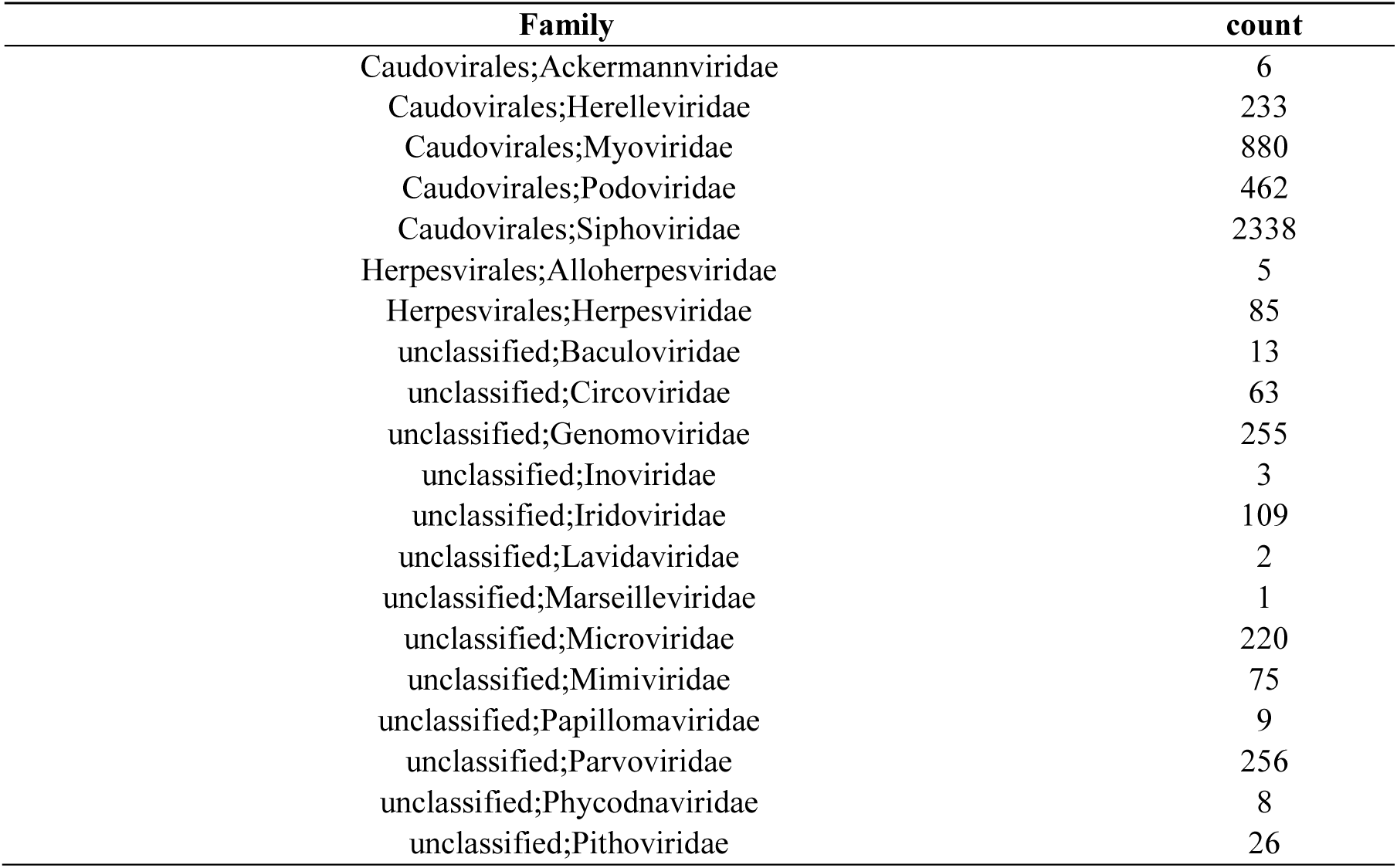
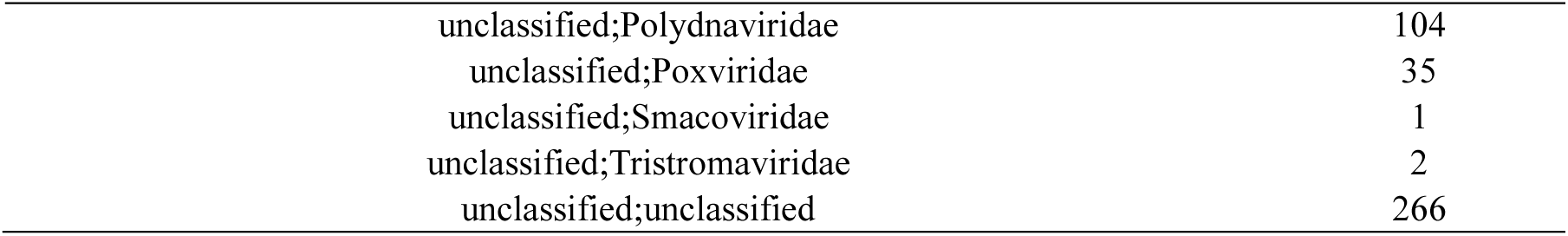
Viral family annotation counts.

Health-dependent restructuring of viral community composition. Marked differences in viral community composition were observed across health states (Table 6). Healthy pangolins harbored more viral families (20–24) compared to diseased (18–20) and deceased individuals (12–16), with a corresponding reduction in genera and species richness in diseased/deceased groups.

**Table 6.** Viral taxonomic richness by health group.

|  | **Healthy MP** | **Healthy MJ** | **Diarrheal MJ** | **Pneumonic MJ** | **Deceased MP** | **Deceased MJ** |
| --- | --- | --- | --- | --- | --- | --- |
| Families | 20 | 24 | 20 | 18 | 16 | 12 |
| Families >1% | 11 | 8 | 7 | 6 | 9 | 3 |
| Genera | 110 | 172 | 105 | 77 | 29 | 25 |
| Genera >1% | 12 | 61 | 12 | 12 | 12 | 10 |
| Species | 475 | 1020 | 440 | 318 | 102 | 83 |
| Species >1% | 29 | 36 | 24 | 15 | 14 | 9 |

MP, Chinese pangolin; MJ, Malayan pangolin.

The most striking difference was in the dominant viral families (Table 7). In all healthy individuals, Caudovirales phages were overwhelmingly dominant: Siphoviridae accounted for 56–59%, followed by Podoviridae (9–18%) and Myoviridae (7–12%). In contrast, diseased and deceased individuals showed dramatic shifts: Parvoviridae accounted for 31% of the pneumonic MJ virome and 66% of the deceased MJ virome; Iridoviridae (38%) and Polydnaviridae (34%) dominated in deceased Chinese pangolins; and Herpesviridae appeared at 9–13% in pneumonic and deceased individuals but was nearly absent in healthy pangolins (<1%).

**Table 7.** Dominant viral families by health group (proportion)

| **Family** | **Healthy MP** | **Healthy MJ** | **Diarrheal MJ** | **Pneumonic MJ** | **Deceased MP** | **Deceased MJ** |
| --- | --- | --- | --- | --- | --- | --- |
| Siphoviridae | 0.56 | 0.59 | 0.68 | 0.49 | - | - |
| Podoviridae | 0.16 | 0.09 | 0.18 | - | - | - |
| Myoviridae | 0.07 | 0.12 | 0.04 | 0.07 | - | - |
| Parvoviridae | - | - | 0.02 | 0.31 | - | 0.66 |
| Iridoviridae | 0.03 | - | - | - | 0.38 | - |
| Polydnviridae | - | - | - | - | 0.34 | - |
| Herpesviridae | 0.01 | - | - | 0.09 | 0.13 | 0.12 |
| Circoviridae | 0.08 | - | - | - | - | - |
| Genoviridae | - | 0.07 | 0.06 | 0.01 | - | - |

"—" indicates proportion < 0.01. Bold values highlight disease-associated expansions.

SIMPER analysis identifies key viruses driving virome dissimilarity. SIMPER analysis quantified the contribution of individual viral taxa to between-group differences (Table 8). At the family level:Diarrheal MJ vs. Healthy MJ: Parvoviridae, Siphoviridae, Myoviridae, and Herpesviridae were the primary contributors.Pneumonic MJ vs. Healthy MJ: Siphoviridae, Podoviridae, Myoviridae, and Genomoviridae were the primary contributors.Deceased MJ vs. Healthy MJ: Parvoviridae, Siphoviridae, and Myoviridae were the primary contributors.Deceased MP vs. Healthy MP: Siphoviridae, Iridoviridae, Polydnaviridae, and Podoviridae were the primary contributors.

**Table 8.** SIMPER analysis: Top contributing viral taxa at family level.

| **Comparison** | **Top contributors (family level)** |
| --- | --- |
| Diarrheal MJ vs. Healthy MJ | Parvoviridae, Siphoviridae, Myoviridae, Herpesviridae |
| Pneumonic MJ vs. Healthy MJ | Siphoviridae, Podoviridae, Myoviridae, Genomoviridae |
| Deceased MJ vs. Healthy MJ | Parvoviridae, Siphoviridae, Myoviridae |
| Deceased MP vs. Healthy MP | Siphoviridae, Iridoviridae, Polydnviridae, Podoviridae |
| Healthy MP vs. Healthy MJ | Siphoviridae, Skunavirus, Podoviridae, Microviridae, Circoviridae |

At the genus level, unclassified Parvoviridae was the top contributor for comparisons involving diseased/deceased Malayan pangolins, while Iridoviridae|Ranavirus and Polydnaviridae|Ichnovirus were the key contributors for Chinese pangolin comparisons. Notably, for the comparison between healthy Chinese and Malayan pangolins, Siphoviridae|Skunavirus, Podoviridae|unclassified, Microviridae|unclassified, and Circoviridae|Cyclovirus were the main discriminators, reflecting species-specific virome signatures.

Alpha and beta diversity. Shannon diversity indices ranged from 2.75 to 7.63 across all 16 samples (Table 9). Although the mean Shannon index was higher in healthy Malayan pangolins than in diarrheal individuals, the difference was not statistically significant (P = 0.11), likely due to limited sample size and high within-group variability.

**Table 9.** Viral alpha diversity (Shannon index)

| Sample | shannon |
| --- | --- |
| SZS1 | 5.88 |
| SZS2 | 7.34 |
| SZS3 | 5.58 |
| SZD1 | 6.63 |
| SZD3 | 2.54 |
| SZD4 | 7.28 |
| SZ15 | 7.63 |
| SZXTY | 6.82 |
| SZXWZ | 6.10 |
| DG241 | 2.75 |
| DG14 | 6.15 |
| DG08 | 4.96 |
| DGB1 | 3.31 |
| DGB2 | 7.33 |
| GZM7 | 5.28 |
| GZM4 | 6.69 |

Hierarchical clustering based on RPKM values revealed clear separation by health status (Figure 1). Deceased and diseased individuals clustered together, diarrheal samples formed a separate branch, and healthy individuals grouped in a distinct cluster, indicating that health status substantially influences viral community structure.

**Figure 1.**
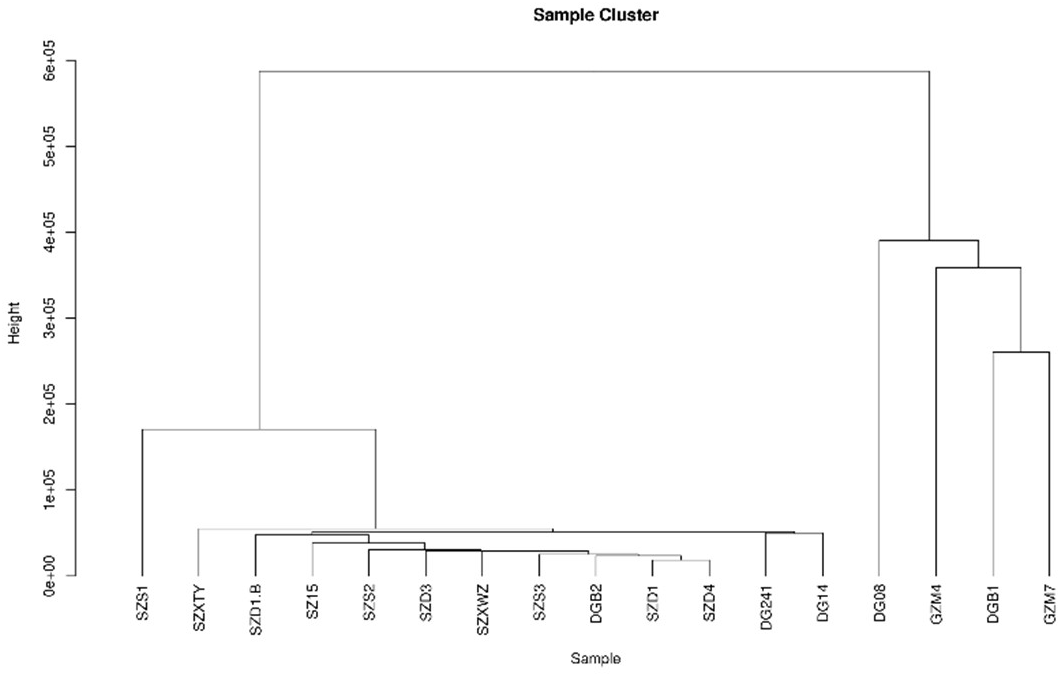
Hierarchical clustering of samples based on viral RPKM values.

Differential biomarker viruses identified by LEfSe. LEfSe analysis identified group-specific biomarker viruses (LDA score > 2.0; Table 10). Free-ranging Malayan pangolins harbored the most biomarker viruses (n = 12), predominantly Staphylococcus phages (Staphylococcus_virus_EW, Staphylococcus_phage_Sebago, Staphylococcus_virus_PH15), Escherichia phages (Escherichia_phage_FEC19, Escherichia_phage_Mansfield, Escherichia_phage_vB_EcoM_WFH), and Bacillus phages (Bacillus_virus_250). Deceased Malayan pangolins had six biomarker viruses, including vertebrate-associated viruses (Alphapapillomavirus_5, Diolcogaster_facetosa_bracovirus, Dasychira_pudibunda_nucleopolyhedrovirus). Healthy Chinese pangolins had six biomarkers, dominated by Streptococcus phages (Streptococcus_phage_phiST1, Streptococcus_phage_Javan414, Streptococcus_phage_Javan235) and Lactobacillus viruses. Notably, no biomarker viruses were identified for captive healthy Malayan pangolins, indicating that their virome composition was the most "average" across all groups.

**Table 10.**
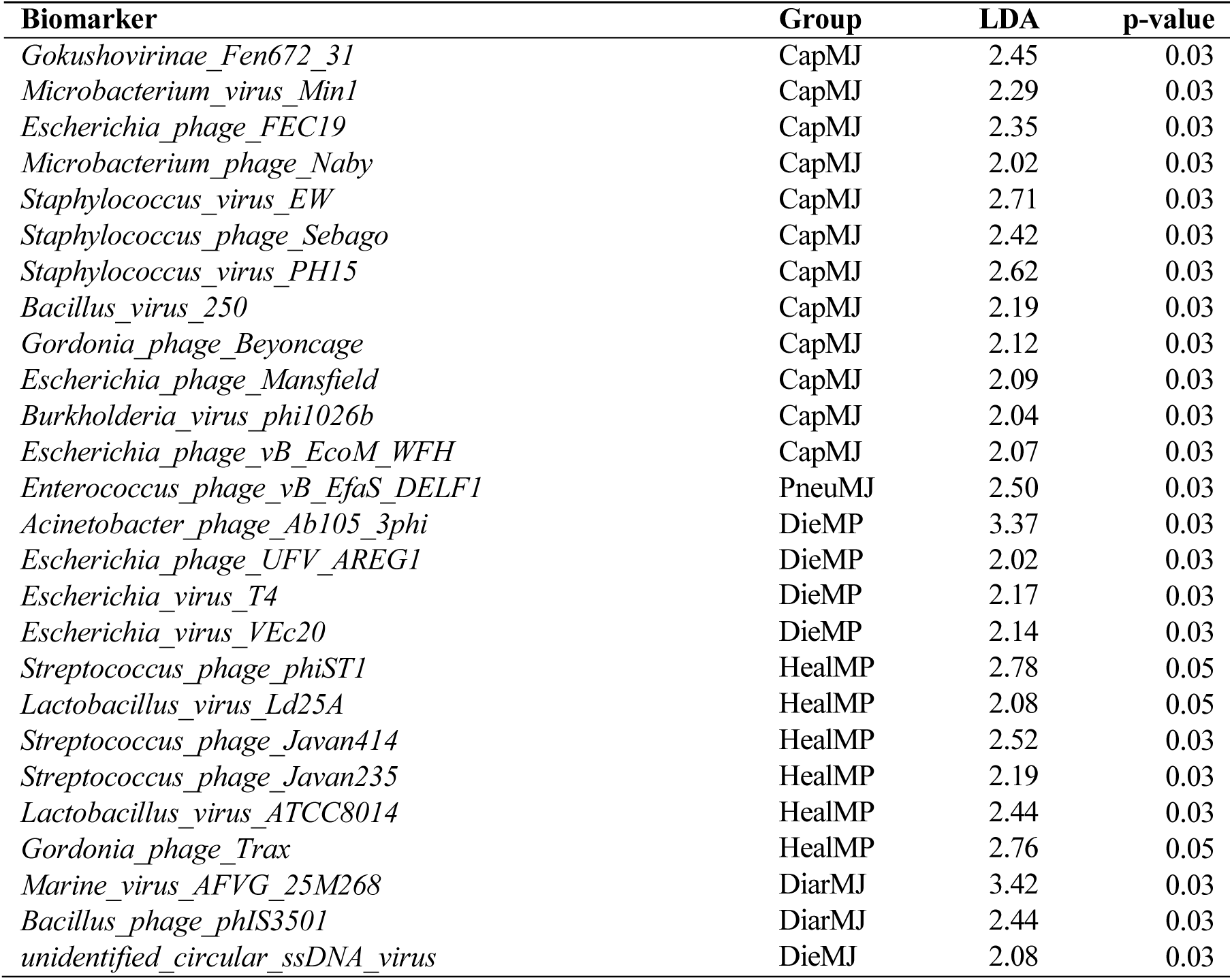
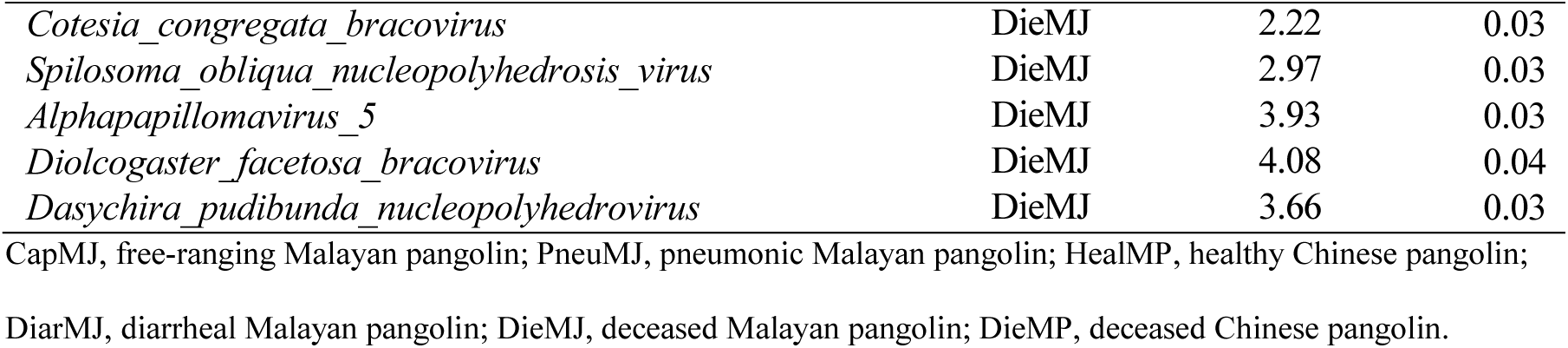
LEfSe biomarker viruses.

Functional annotation of the pangolin gut virome. KEGG functional annotation at Level 3 identified 121 functional categories. The most gene-rich functions were DNA repair and recombination proteins (ko03400; 300 genes), DNA replication proteins (ko03032; 239 genes), DNA replication (ko03030; 162 genes), pyrimidine metabolism (ko00240; 127 genes), mismatch repair (ko03430; 123 genes), homologous recombination (ko03440; 119 genes), prokaryotic defense systems (ko02048; 102 genes), purine metabolism (ko00230; 68 genes), and chromosome and associated proteins (ko03036; 82 genes) (Figure 2). The predominance of genes related to DNA replication, repair, and recombination indicates that the pangolin gut virome undergoes frequent genomic replication and recombination events.

**Figure 2.**
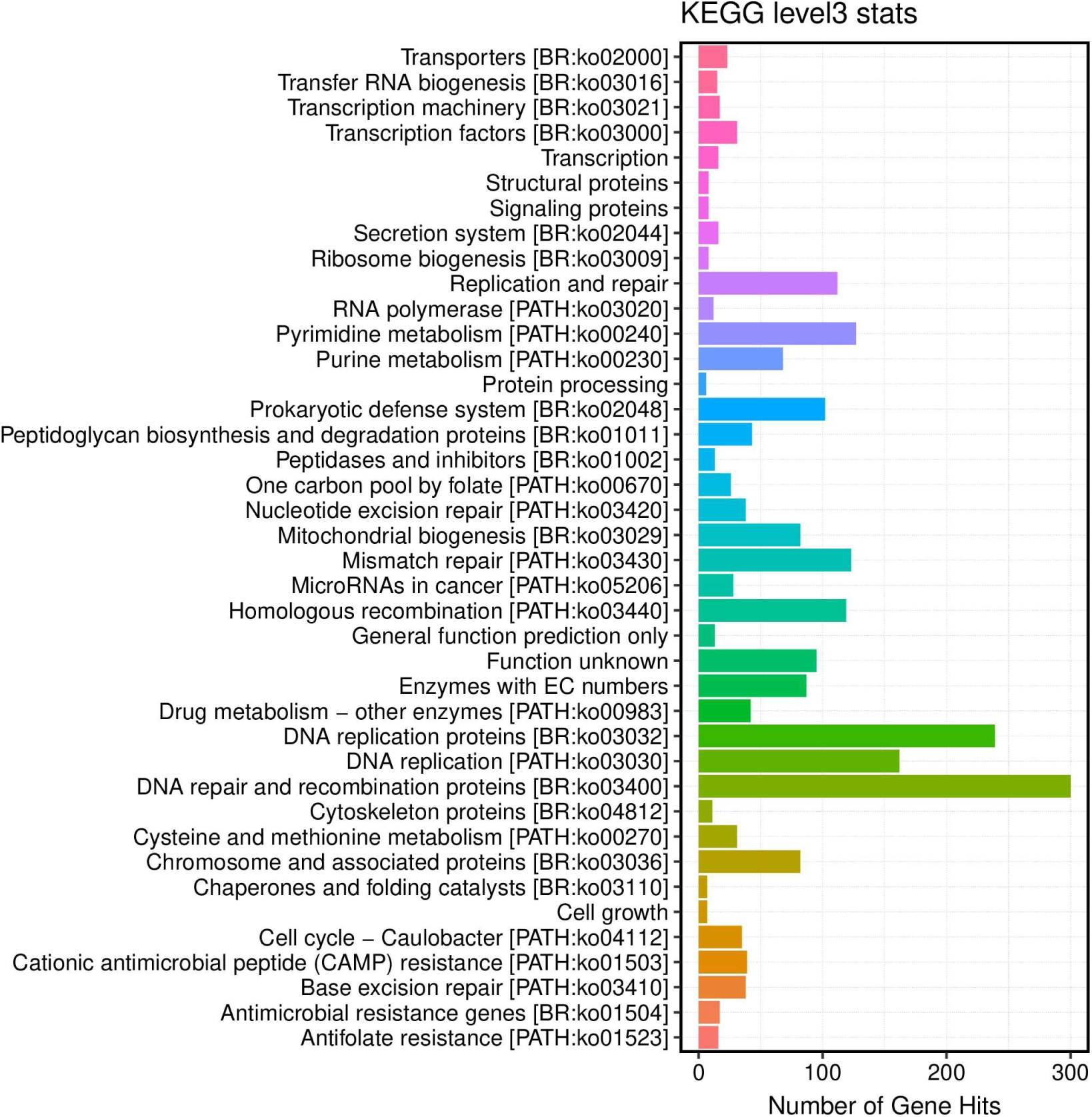
KEGG Level 3 functional annotation of the pangolin gut virome.

## Discussion

This study provides the first comprehensive characterization of the gut DNA virome in pangolins across multiple health states. Our findings reveal that the pangolin gut virome is dominated by Caudovirales phages in healthy individuals but undergoes a fundamental restructuring during disease, characterized by phage depletion and eukaryotic virus expansion.

Phage depletion as a hallmark of intestinal virome dysbiosis. The most striking finding of this study is the dramatic reduction in phage content in diseased and deceased pangolins compared to healthy individuals (94.5% vs. 53.3%, P = 0.02). This pattern parallels observations in COVID-19 patients, where non-COVID-19 feces were enriched in phages while COVID-19 patients showed increased eukaryotic viruses alongside Escherichia and Enterobacter phages (11, 12). However, the direction of phage change appears disease-dependent: in Clostridioides difficile infection (CDI), Caudovirales phage abundance was significantly increased compared to healthy controls (13), and in type 2 diabetes (T2D), Myoviridae, Podoviridae, and Siphoviridae relative abundances were increased (19). These contrasting patterns suggest that phage responses to disease are context-dependent, likely influenced by the nature of the insult (viral vs. bacterial vs. metabolic), the host immune status, and the baseline virome composition. In pangolins, the consistent depletion of phages across multiple disease states (diarrhea, pneumonia, and death) suggests a generalizable mechanism—possibly involving host immune suppression allowing opportunistic eukaryotic viral expansion, or disease-induced disruption of bacterial communities that sustain phage populations.

Parvoviridae and Iridoviridae as candidate pathogenic viruses. SIMPER analysis consistently identified Parvoviridae as the primary contributor to virome dissimilarity between healthy and diseased/deceased Malayan pangolins. Parvoviridae accounted for only 2% of the diarrheal virome but expanded to 31% in pneumonic and 66% in deceased Malayan pangolins. Parvoviruses are small, non-enveloped ssDNA viruses known to cause severe gastrointestinal disease in multiple mammalian species, including canine parvovirus (CPV) and porcine parvovirus (PPV) (33). The dramatic expansion of Parvoviridae in diseased pangolins suggests either direct pathogenic invasion or opportunistic expansion following immune compromise. Further studies using PCR quantification and histopathological examination are needed to establish causality.

For Chinese pangolins, Iridoviridae (38%) and Polydnaviridae (34%) were the dominant viruses in the deceased individual. Iridoviruses, particularly Ranavirus species, are known primarily as pathogens of amphibians, reptiles, and fish (34), but have been increasingly detected in mammals (35). The detection of Ranavirus-like sequences in deceased Chinese pangolins raises concerns about potential cross-class transmission, possibly facilitated by the insectivorous diet of pangolins. Polydnaviridae, which are typically associated with parasitoid wasps, may represent dietary or environmental contamination rather than true infection; however, their high relative abundance warrants further investigation.

Species-specific virome signatures. Healthy Chinese pangolins harbored six biomarker viruses (primarily Streptococcus phages and Lactobacillus viruses), while no biomarker viruses were identified for captive healthy Malayan pangolins. This difference likely reflects the more diverse virome composition of Chinese pangolins, which may compensate for their lower gut bacterial functional diversity. The observation that Chinese pangolins possess higher viral diversity but lower bacterial functional diversity than Malayan pangolins suggests a complementary relationship between the bacterial and viral components of the gut microbiome, where virome complexity may partially buffer the effects of reduced bacterial functional capacity.

Free-ranging Malayan pangolins had the highest number of biomarker viruses (n = 12), predominantly Staphylococcus and Escherichia phages, likely reflecting exposure to a broader range of environmental microbes and viruses compared to captive individuals. This finding supports the hypothesis that environmental contact enriches virome diversity, which may contribute to immune priming and resilience (36).

Frequent genomic recombination in the pangolin gut virome. KEGG functional annotation revealed that genes related to DNA replication, repair, and recombination were the most abundant in the pangolin gut virome, followed by purine and pyrimidine metabolism. This functional profile is consistent with active viral replication and frequent homologous recombination events, which are hallmarks of phage evolution in gut ecosystems (37). The high abundance of prokaryotic defense system genes (ko02048; 102 genes) further suggests an ongoing arms race between phages and their bacterial hosts, driving rapid co-evolution and genomic diversification.

Limitations. Several limitations should be acknowledged. First, the sample size for some groups (particularly deceased and pneumonic individuals) was small, limiting statistical power. Second, this study focused exclusively on DNA viruses; RNA viruses (including potentially important pathogens such as coronaviruses) were not captured by our sequencing approach. Third, the cross-sectional design precludes causal inference—whether viral changes precede or follow disease onset cannot be determined without longitudinal sampling. Fourth, the detection of Parvoviridae and Iridoviridae by metagenomic sequencing requires validation by targeted PCR and isolation studies to confirm active infection rather than passive passage of viral DNA.

Conclusions. This study demonstrates that the gut virome of pangolins is closely associated with health status. Healthy pangolins harbor a phage-dominated virome with Caudovirales as the principal component, whereas disease triggers a shift toward eukaryotic virus dominance. Parvoviridae emerges as the key virus associated with disease in Malayan pangolins, while Iridoviridae is the primary virus in deceased Chinese pangolins. These findings have important implications for pangolin conservation: (1) parvovirus surveillance should be implemented in rescue centers as a potential health indicator; (2) phage content may serve as a biomarker of gut virome health; and (3) free-ranging environments appear to promote virome diversity, supporting the creation of semi-natural enclosures for rescued pangolins. Future studies incorporating RNA viromics, longitudinal sampling, and experimental validation of candidate pathogenic viruses will further advance our understanding of the pangolin gut virome and its role in health and disease.

## Materials and Methods

Sample collection. Fecal samples were collected from pangolins at the Shenzhen Wildlife Rescue Center and Dongguan rescue facility, China. In October 2020, 10 samples were collected from two adult Chinese pangolins, six adult Malayan pangolins, and two subadult Malayan pangolins. In September 2021, 16 samples were collected from individuals across six health categories: diarrheal Malayan pangolins (n = 3), pneumonic Malayan pangolins (n = 2), healthy Chinese pangolins (n = 2), free-ranging Malayan pangolins (n = 2), healthy Malayan pangolins (n = 5), and deceased pangolins (n = 2). Fresh feces were collected within 2 h of defecation and immediately stored at −80°C until processing. Detailed sample information is provided in Table 1.

Virus-like particle (VLP) enrichment and nucleic acid extraction. Samples were homogenized and subjected to low-speed centrifugation to remove tissue debris and cells. VLPs were purified and concentrated through a combination of filtration (0.45 μm membrane) and ultracentrifugation. Viral nucleic acids (dsDNA, ssDNA, ssRNA, and dsRNA) were co-extracted from purified VLPs. For ssDNA viruses, complementary strands were synthesized prior to library construction. DNA libraries were prepared by random fragmentation (ultrasonication), end repair, adapter ligation, and PCR amplification. Qualified libraries were sequenced on the Illumina platform (PE150).

Quality control and host removal. Raw reads were quality-filtered using Trimmomatic (25). Clean reads were aligned to the host reference genomes (Manis javanica and M. pentadactyla) using BWA v0.7.17 (mem -k 30) (26). Reads mapping to host genomes with ≥80% alignment length was removed.

De novo assembly and contig clustering. Host-filtered clean reads were assembled using Megahit v1.1.2 (--presets meta-large --min-contig-len 300) (27). Assembled contigs were remapped with BWA to calculate assembly rates. Residual host contigs were identified by BLAST (v2.9.0+) against host genomes and removed (criteria: alignment length ≥500 bp and similarity ≥90%, or alignment covering ≥80% of contig length with similarity ≥90%). All contigs were clustered using CD-HIT v4.7 (-c 0.95 -aS 0.8) (28) to obtain non-redundant unique contigs.

Viral sequence identification. A two-strategy approach was employed to maximize viral detection while minimizing false positives.

Strategy 1: Reference-based identification. Unique contigs were compared against the Virus-NT database using BLAST (v2.9.0+). Contigs meeting similarity ≥80%, alignment length ≥500 bp, and e-value ≤1e-5 were classified as "confirmed"; those with alignment length ≥100 bp and e-value ≤1e-5 but not meeting the confirmed criteria were classified as "suspected."

Strategy 2: De novo identification (29, 30). Candidate viral sequences were identified if they met any of three criteria: (A) blastn against Virus-NT (e ≤1e-5); (B) blastx against Virus-NR (e ≤1e-3); or (C) MetaGeneMark v3.38 gene prediction followed by hmmsearch v3.2.1 against VPFs and vFam HMM databases (e ≤1e-5). False positives were excluded by: (A) blastn against the full NT database (e ≤1e-10); (B) diamond v0.9.10 against NR (e ≤1e-3); and (C) NCBI Taxonomy annotation—sequences with ≥20% of top 50 BLAST hits annotated as non-viral (Eukaryota, Bacteria, or Archaea) were excluded. Final annotations were based on best-hit results against Virus-NT (length ≥100 bp, e ≤1e-5).

Viral abundance calculation. Viral abundance was calculated as RPKM (Reads Per Kilobase per Million mapped reads): RPKM = (Contig reads) / (Total mapped reads / 10^6 × Contig length / 10^3).

Alpha and beta diversity analysis. Shannon diversity indices were calculated based on RPKM values for each sample. Bray-Curtis distances were computed and hierarchical clustering (hclust) was performed to assess beta diversity. Group comparisons for alpha diversity were conducted using Wilcoxon rank-sum tests.

Differential biomarker analysis. LEfSe (Linear Discriminant Analysis Effect Size) was used to identify viral taxa significantly associated with specific health groups (31). An LDA score threshold of 2.0 was applied. SIMPER (Similarity Percentage) analysis was performed using the R package vegan to quantify the contribution of individual viral taxa to between-group dissimilarities.

Functional annotation. Non-redundant gene sequences were compared against the KEGG gene database using diamond (32). Functional pathways were annotated at KEGG Level 3.

Statistical analysis. One-way ANOVA was used to compare phage proportions between healthy and diseased/deceased groups. All statistical analyses were performed in R v4.0. P-values < 0.05 were considered significant.

Accession number(s). Raw sequencing data have been deposited in NCBI SRA under BioProject accession number [PRJNA1014131 to be assigned upon submission]. Assembled contigs and annotation files are available from the corresponding author upon reasonable request.

## Acknowledgments

We thank the Shenzhen Wildlife Rescue Center and Dongguan rescue facility for providing samples.

## Author contributions

W.J. designed the study, performed experiments, analyzed data, and wrote the manuscript. J.P. supervised the project and revised the manuscript. Both authors read and approved the final manuscript.

## Funding

This work was supported by GDAS Special Project of Science and Technology Development (2022GDASZH-2022010106) and Planning Funds of Scientific and Technology of Guangdong Province (No. 2023A1111110001 and 2024B1212040009).

## Competing interests

The authors declare no competing interests.

The procedures for care and use of animals were approved by the Ethics Committee of the Institute of Zoology, Guangdong Academy of Sciences (Guangzhou, China) and all applicable institutional and governmental regulations concerning the ethical use of animals were followed.

## Availability of data and materials

Raw sequencing data have been deposited in NCBI SRA under BioProject accession number [PRJNA1014131 to be assigned upon submission]. Assembled contigs and annotation files are available from the corresponding author upon reasonable request.

## Consent for publication

Not applicable.

